# Evaluating the neural underpinnings of motivation for walking exercise

**DOI:** 10.1101/2022.12.30.522346

**Authors:** Sarah Doren, Sarah M. Schwab, Kaitlyn Bigner, Jenna Calvelage, Katie Preston, Abigail Laughlin, Colin Drury, Brady Tincher, Daniel Carl, Oluwole O. Awosika, Pierce Boyne

## Abstract

**Background:** Motivation is critically important for rehabilitation, exercise, and motor performance, but its neural basis is poorly understood. Recent correlational research suggests that superior frontal gyrus medial area 9 (SFG9m) may be involved in motivation for walking activity. This study experimentally evaluated brain activity changes in periods of additional motivation during walking exercise, and tested how these brain activity changes relate to self-reported exercise motivation and walking speed.

**Methods:** Non-disabled adults (N=26; 65% female; 25 ± 5 years old) performed a vigorous exercise experiment involving 20 trials of maximal speed overground walking. Half of the trials were randomized to include ‘extra motivation’ stimuli (lap timer, tracked best lap time and verbal encouragement). Wearable nearinfrared spectroscopy measured oxygenated hemoglobin responses (ΔHbO_2_) from frontal lobe regions, including the SFG9m, primary motor, dorsolateral prefrontal, anterior prefrontal, supplementary motor and dorsal premotor cortices.

**Results:** Compared with standard trials, participants walked faster during ‘extra-motivation’ trials (2.67 vs. 2.43 m/s; p<0.0001) and had higher ΔHbO_2_ in all tested brain regions. This extra motivation effect on ΔHbO_2_ was greatest for SFG9m (+703 µM) compared with other regions (+45 to +354 µM; p≤0.04). Greater SFG9m activity was correlated with more self-determined motivation for exercise and faster walking speed.

**Conclusions:** Simple motivational stimuli during walking exercise seem to upregulate widespread brain regions, especially SFG9m. This could help explain the positive effects of motivational feedback on gait outcomes observed in prior rehabilitation research. Thus, these findings provide a potential biologic basis for the benefits of motivational stimuli, elicited with clinically-feasible methods during walking exercise. Future clinical studies could build on this information to develop prognostic biomarkers and test novel brain stimulation targets for enhancing exercise motivation (e.g. SFG9m).

## Introduction

Human motor behavior emerges from the interaction between environmental and individual constraints that are further bounded by a task goal.^1-7^ Motivation is a critical constraint to consider in explaining how individuals approach a movement task, are pushed to discover novel task solutions, and ultimately improve motor performance with practice.^7^ Self-determination theory^8-10^ differentiates between intrinsic and extrinsic forms of motivation, where *intrinsic motivation* refers to performing an action because it is inherently satisfying.^8-11^ *Extrinsic motivation*, on the other hand, refers to actions driven by external pressures or rewards.^9^

Many human behaviors begin with extrinsic motivation, then become more *self-determined* as individuals progress from strictly *external* regulation to higher degrees of *introjection* (seeking approval from self or others), then *identification* (conscious valuing of activity), and eventually, *intrinsic* motivation.^9^ Behaviors regulated primarily by external and introjected motivations are considered to be *controlled*, whereas behaviors regulated primarily by identified and intrinsic motivations are considered *autonomous*.^10^

Motivation for physical activity, especially in its more self-determined forms,^12, 13^ can play an important positive role in an individual’s engagement, physical performance, motor skill acquisition, and outcomes in physical rehabilitation.^14-19^ For example, greater exercise motivation during inpatient stroke rehabilitation has been associated with increased participation in physical activity after discharge (especially intrinsic, identified and introjected motivation).^13^ Daily feedback about self-selected fast gait speed during inpatient stroke rehabilitation has also been shown to significantly improve gait speed at discharge,^20^ potentially by augmenting identified motivation (assuming participants valued faster walking), or even intrinsic motivation (e.g. if it made rehabilitation feel more like a game).

While motivation for physical activity is generally considered to be critically important for rehabilitation, exercise, and motor performance, it has largely been studied using subjective questionnaires or interviews (e.g.^13, 21-24^), and the neural basis remains poorly understood. A better understanding of the neurobiology of exercise motivation could lead to: more objective measurement (including unconscious aspects not captured by questionnaires); more accurate prognostication of movement recovery after injury; better prediction of response to motor rehabilitation or exercise interventions; and novel brain stimulation targets to enhance motivation for exercise. Broadly speaking, different aspects of motivation have been related to interactions among various brain regions, including the dorsolateral prefrontal cortex (DLPFC), anterior prefrontal cortex (aPFC), supplementary motor area (SMA), anterior cingulate cortex (ACC), ventral striatum, and anterior insula.^25-27^

However, technical challenges have made it difficult to measure motivation-related brain activity *during* mobility tasks like walking that involve whole-body transport and continuous head motion.^28^ Thus, prior studies aiming to probe the neural basis of motivation for mobility activities have primarily been limited to correlational designs. Intriguingly, these correlational mobility studies have suggested a possible motivation-related role for a previously unconsidered brain region called superior frontal gyrus medial area 9 (SFG9m),^28, 29^ which is located in the medial prefrontal cortex just superficial to the ACC. Non-disabled adults who had greater SFG9m resting connectivity during a brain MRI also walked further during a 6-minute walk test,^28^ and persons with and without stroke who had greater SFG9m activation during imagined walking inside an MRI scanner also had better overall gait function.^29^ Still, the asynchronous measurement of brain activity and walking function in these correlational studies makes it impossible to draw any conclusions about the role of SFG9m or any other brain regions in motivation for walking activity.

Functional near-infrared spectroscopy (fNIRS) may help overcome this technical limitation by enabling the study of brain activity *during* mobility tasks like walking. fNIRS uses infrared light to measure changes in blood oxygenation, which can be used to infer cerebral cortex activity when the light sources and detectors are arranged on the scalp.^30^ Unlike other imaging modalities, head motion does not create signal artifact with fNIRS, as long as the light sources and detectors maintain their scalp positions.^30^ Thus, fNIRS is well suited to studying mobility tasks. However, no prior walking studies have capitalized on fNIRS to experimentally assess differences in brain activity underlying different motivational conditions.

The purpose of this fNIRS study was to assess brain activity changes in periods of additional motivation during walking exercise. We also tested how the magnitude of these brain activity changes relates to self-reported motivation for exercise, and to differences in walking speed between trials. In line with previous correlational studies,^28, 29^ we hypothesized that SFG9m would be more active during periods of additional motivation and would have greater motivation-related activity than other brain regions. We also hypothesized that greater SFG9m activation would be associated with more self-determined motivation and faster walking.

## Methods

This study was approved by the University of Cincinnati Institutional Review Board and performed between February 2021 and March 2022. Non-disabled adult participants were recruited from the University community to complete online questionnaires and perform a single visit walking experiment with fNIRS. Written informed consent was obtained prior to participation.

### Participants

To be eligible, participants had to be 18 to 80 years old, able to communicate with investigators and provide informed consent, not pregnant, and without any health conditions that would require medical clearance for vigorous exercise according to the American College of Sports Medicine preparticipation screening algorithm.^31^

### Experimental Design

Each participant performed 20 total trials of maximal speed overground walking back and forth along a straight 12-meter course while wearing the fNIRS system (Fig 1A-B). Each trial included two laps back and forth (48 meters). Between trials, there was a mandatory 20-second standing rest break, then participants were able to start the next trial when ready. A computer was placed on a raised table at the starting and finishing end of the course, while floor tape marked the other end. ‘PyschoPy’ software^32^ was used to present audiovisual stimuli, randomize trial conditions, time laps and wirelessly transmit timestamps for each trial to the fNIRS data collection software, using the lab streaming layer protocol. Participants pressed a large button connected to the computer to start each trial, start the second lap and finish the trial.

**Figure 1.**
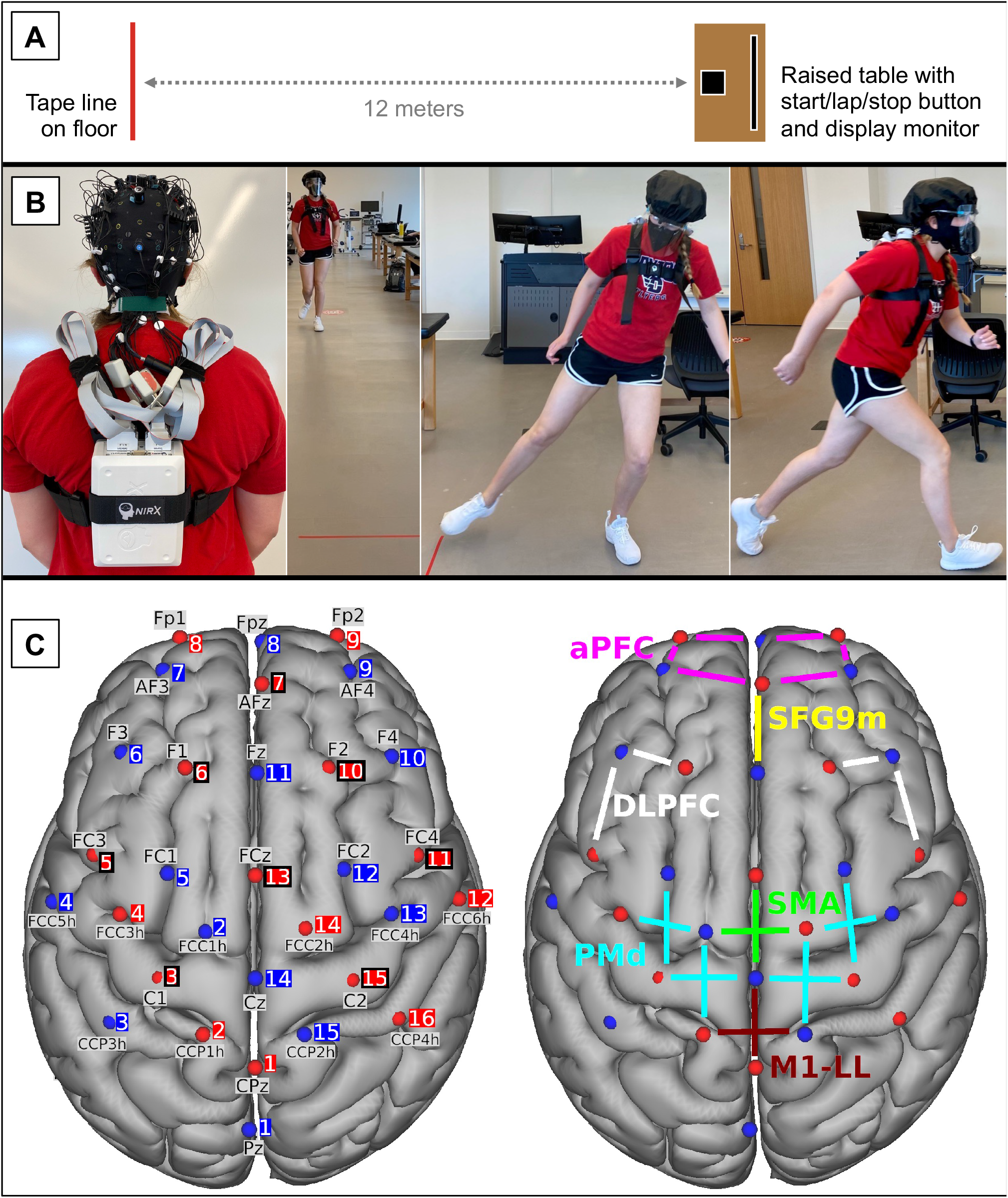
Experimental setup. (A) Walking course schematic. (B) fNIRS cap setup and wire management (left) for trial performance (right). (C) Locations of fNIRS sources (red spheres) and detectors (blue spheres) are visualized on the cortical surface of the MNI152 standard brain model. Left panel: Each optode is shown with its standard 10/5% location label and source or detector number. Black squares around source numbers 3, 5, 6, 7, 10, 11, 13 & 15 show the locations of the 8 short-channel detectors. Right panel: Lines between sources and detectors represent channels that were averaged to represent each color-coded region of interest. fNIRS, functional near-infrared spectroscopy; SFG9m, superior frontal gyrus medial area 9; M1-LL, primary motor cortex lower limb area; DLPFC, dorsolateral prefrontal cortex; aPFC, anterior prefrontal cortex; SMA, supplementary motor area; PMd, dorsal premotor cortex.

Participants were instructed to walk as fast as possible (without running) for all trials, and half of the trials were randomized to include additional audiovisual stimuli intended to augment motivation. For these ‘extra motivation’ trials, the computer displayed a large lap timer and the participant’s current best lap time, provided an auditory count of the lap time every five seconds, and said “new record!” whenever the best lap time was updated. Throughout the extra motivation trials, at least two study team members also provided verbal encouragement. None of these extra stimuli were presented during standard trials.

Throughout the experiment, heart rate was also recorded at 0.33 Hz using an electrocardiographic monitor (Polar H7-H10), and participants were required to wear a face mask due to the COVID-19 pandemic.

### fNIRS Data Collection

A wearable fNIRS system (NIRSport2, NIRx Medical Technologies) was used to record brain activity during the experiment (Fig 1B) at a sampling rate of 5.09 Hz. The 32 dual-tip optodes included 16 light sources emitting infrared wavelengths of 760 and 850 nm, and 16 silicon photodiode detectors. Optodes were secured to the head at landmarks from the 10-5% EEG location system^33^ using an EasyCap (EASYCAP GmbH, Germany) and spring-loaded optode holders. Wires were managed with cable trees, ties, and holders to avoid tension on the optodes during walking, and a black shower cap was placed over the optodes to block environmental light.

Brain regions of interest included SFG9m, primary motor cortex lower limb area (M1-LL), DLPFC, aPFC, SMA and dorsal premotor cortex (PMd). The optode configuration (Fig 1C, Table 1) was designed by visualizing the Human Connectome Project cortical atlas^34^ alongside the average locations of the 10-5% landmarks in the same standard MNI brain space.^35^ Source-detector pairs with approximately 3 cm separation formed standard channels to record cortical activity from the regions of interest. Eight short-distance detectors with 8 mm source separation also formed short-distance channels to record signal of non-interest superficial to the brain. This can be used to filter motion artifacts (from cap slippage on the head) and global physiologic noise (e.g. from hemodynamic fluctuations) out of the standard channel data.^36-39^ The short-distance detector bundle occupied one of the 16 detector optodes.

**Table 1.** fNIRS Optode Configuration.

| Table 1. fNIRS Optode Configuration |  |  |  |  |  |
| --- | --- | --- | --- | --- | --- |
| Source | 10-5% Label | MNI Coordinates (mm) | Detector | 10-5% Label | MNI Coordinates (mm) |
| **S1** | CPz | 0.83, -41.89, 75.59 | **D1** | Pz | -0.98, -66.75, 62.61 |
| **S2** | CCP1h | -15.04, -30.55, 77.26 | **D2** | FCC1h | -14.19, 1.2, 72.89 |
| **S3** | C1 | -28.43, -14.43, 71.27 | **D3** | CCP3h | -43.19, -30.52, 65.78 |
| **S4** | FCC3h | -40.33, 2.49, 60.71 | **D4** | FCC5h | -60.8, -1.96, 37.33 |
| **S5** | FC3 | -48.24, 16.31, 45.55 | **D5** | FC1 | -25.88, 16.41, 62.65 |
| **S6** | F1 | -20.49, 44.82, 46.59 | **D6** | F3 | -39.81, 44.33, 31.74 |
| **S7** | AFz | 2.90, 64.49, 25.87 | **D7** | AF3 | -27.23, 64.88, 14.87 |
| **S8** | Fp1 | -22.08, 67.58, -6.47 | **D8** | Fpz | 3.02, 67.61, -3.04 |
| **S9** | Fp2 | 25.86, 68.06, -7.13 | **D9** | AF4 | 29.57, 64.55, 15.08 |
| **S10** | F2 | 22.99, 44.91, 47.06 | **D10** | F4 | 42.36, 43.57, 33.13 |
| **S11** | FC4 | 50.05, 16.47, 46.51 | **D11** | Fz | 1.46, 44.16, 49.44 |
| **S12** | FCC6h | 62.88, -1.1, 37.31 | **D12** | FC2 | 27.78, 17.88, 63.08 |
| **S13** | FCz | 0.91, 16.38, 64.36 | **D13** | FCC4h | 42.27, 2.62, 60.65 |
| **S14** | FCC2h | 16.03, 2.4, 73.06 | **D14** | Cz | 0.88, -13.47, 74.14 |
| **S15** | C2 | 30.3, -14.79, 71.79 | **D15** | CCP2h | 15.81, -30.26, 77.98 |
| **S16** | CCP4h | 44.43, -29.43, 65.85 |  |  |  |
| Brain Region |  |  | Channels |  |  |
| Superior frontal gyrus medial area 9 (SFG9m) |  |  | S7-D11 |  |  |
| Primary motor cortex lower limb area (M1-LL) |  |  | S1-D14, S2-D15 |  |  |
| Dorsolateral prefrontal cortex (DLPFC) |  |  | S5-D6, S6-D6, S10-D10, S11-D10 |  |  |
| Anterior prefrontal cortex (aPFC) |  |  | S7-D7, S7-D9, S8-D7, S8-D8, S9-D8, S9-D9 |  |  |
| Supplementary motor area (SMA) |  |  | S13-D14, S14-D2 |  |  |
| Dorsal premotor cortex (PMd) |  |  | S2-D2, S3-D14, S3-D5, S4-D2, S14-D15, S14-D13, S15-D14, S15-D12 |  |  |

### fNIRS Experiment Data Processing

The NIRS Toolbox^40^ (version 837) and a custom Matlab script were used to process data. fNIRS voltage data were down sampled to 5 Hz and converted to optical densities then hemoglobin concentrations using the Beer-Lambert Law.^40, 41^ The first 6 principal components from the short-channel data were regressed out of the standard channel data using autoregressive, iteratively-reweighted least squares regression.^39, 40, 42^ The filtered oxygenated hemoglobin (HbO_2_) data were then averaged among channels representing the same cortical region (Fig 1C).

The first two (practice) trials were discarded, leaving 18 for analysis (9 extra motivation and 9 standard trials). Since walking speed and trial duration varied between trials, the time scale was normalized by interpolating HbO_2_ data to percentage of trial duration. For each cortical region and each trial, HbO_2_ response (ΔHbO_2_) was calculated as the mean HbO_2_ value from 0-110% trial duration, minus the mean HbO_2_ value just before trial onset (from -20% to 0% trial duration).

HR data were expressed as a percentage of age-predicted maximal HR (AP-HR_max_), which was calculated using 206.9 – 0.67 x age (years).^43^ HR was averaged across the whole session to measure overall exercise intensity, and was averaged from 100-150% trial duration to capture the response for each trial.

### Self-Report Measures

The following questionnaires were completed online within 2 ± 3 days (mean ± SD) of the fNIRS experiment:

#### The International Physical Activity Questionnaire (IPAQ)^44^

short form is a 9-item questionnaire that collects information on the time spent participating in vigorous-intensity, moderate-intensity, walking and sitting activities over the last 7 days. Results were used for descriptive purposes to estimate total metabolic equivalents (METs) of energy expenditure per week and to categorize individuals as having low, moderate or high amounts of physical activity.^44^

#### The Behavioral Regulation in Exercise Questionnaire (BREQ)^45^

includes 15 items assessing motivation for exercise, based on self-determination theory. Each item is rated from zero (not true for me) to four (very true for me). Subsets of questions were averaged to calculate scores for different dimensions of motivation (external, introjected, identified, and intrinsic),^9, 46^ and subsets of these domains were averaged to calculate scores for autonomous (intrinsic and identified) and controlled (introjected and external) motivations.^47^ The primary measure of self-determined motivation for exercise was the difference between the autonomous and controlled motivation scores.^47^

### Statistical Analysis

SAS software (SAS Institute, Cary, NC) version 9.4 was used for analysis, except as indicated. The effects of trial condition (extra motivation vs. standard) on walking speed, heart rate and brain ΔHbO_2_ were assessed using separate linear regression models. The models included data from each trial and assumed compound symmetry covariance between repeated trials from the same participant.

In the brain ΔHbO_2_ (primary) model, condition effects and (co)variances were estimated separately for each brain region, and trial HR was included as a covariate. This covariate was added to control for potential confounding effects of global hemodynamic differences (i.e. higher HR) on brain ΔHbO_2_ differences between conditions, in case the short-channel regression did not fully remove such physiologic noise. Within this model, the effect of extra motivation on ΔHbO_2_ was estimated for each brain region by contrasting the two conditions, and the magnitude of this extra motivation effect for SFG9m (the primary region of interest) was compared to each other region.

We also assessed how individual differences in the brain responses to extra motivation related to self-reported motivation for exercise. For this analysis, mean extra motivation effects on ΔHbO_2_ for each participant and brain region were calculated by averaging across trials within conditions. Partial Pearson correlations were then calculated between these brain responses and BREQ scores, controlling for each participant’s mean HR across all trials.

To assess how brain activity related to walking speed, we used data from each trial to calculate partial repeated measures correlations between brain ΔHbO_2_ and speed, controlling for trial condition (extra motivation vs. standard). Repeated measures correlations estimate common within-participant associations,^48^ and were calculated using R software^49^ with the ‘rmcorr’ package.^48^

### Sample Size

Power analysis was done using the R package ‘pwr’.^50^ Simplified calculations based on a paired t-test indicated that 18 participants would provide 80% power at the two-sided 0.05 significance level to detect a medium standardized effect (Cohen’s d of 0.5) between the extra motivation and standard conditions in the primary analysis. For the correlation analyses, simplified calculations based on a bivariate Pearson correlation without repeated measures indicated that 23 participants would provide 80% power to detect a correlation as small as 0.55. Thus, we aimed to enroll at least 23 participants.

## Results

Among 30 persons who consented to participate, four were ultimately unavailable to perform the fNIRS experiment. The remaining 26 participants (Table 2) were enrolled and completed the study, including the full fNIRS experiment and both self-report measures.

**Table 2.** Participant Characteristics (N=26)

| **Table 2. Participant Characteristics (N=26)** |  |
| --- | --- |
| Age, years | 24.8 ± 5.0 [18.8, 42.1] |
| Female, n (%) | 17 (65.4%) |
| Body mass index, kg/m<sup>2</sup> | 24.8 ± 3.9 [19.2, 35.3] |
| IPAQ total physical activity, MET-minutes/week | 2560 ± 1273 [330, 5346] |
| IPAQ weekly physical activity category |  |
| High | 13 (50.0%) |
| Moderate | 12 (46.2%) |
| Low | 1 (3.8%) |
| BREQ motivation for exercise scores |  |
| Overall self-determined motivation, -4 to 4<br>(Autonomous – Controlled) | 0.0 ± 0.6 [-1.1, 1.2] |
| Autonomous motivation, 0 to 4<br>mean (Intrinsic, Identified) | 2.4 ± 0.6 [0.9, 3.3] |
| Intrinsic motivation, 0 to 4 | 2.8 ± 0.5 [1.0, 4.0] |
| Identified motivation, 0 to 4 | 2.0 ± 0.6 [0.5, 3.0] |
| Controlled motivation, 0 to 4<br>mean (Introjected, External) | 2.4 ± 0.5 [1.7, 3.4] |
| Introjected motivation, 0 to 4 | 2.1 ± 0.5 [1.3, 3.0] |
| External motivation, 0 to 4 | 2.7 ± 0.6 [1.3, 3.8] |
| Values are mean ± SD [range] or N (%). IPAQ, International Physical Activity Questionnaire; MET, metabolic equivalents of energy expenditure; BREQ, Behavioral Regulation in Exercise Questionnaire |  |

### Walking Exercise Intensity

The average HR across the fNIRS experiment was 84.7 ± 8.1 % AP-HR_max_ (mean ± SD across participants). Compared with standard trials, participants walked faster during extra motivation trials (2.67 ± 0.32 vs. 2.43 ± 0.31 m/s; p<0.0001) and had slightly higher heart rates (89.1 ± 8.5 vs. 87.8 ± 8.3 % AP-HR_max_, p<0.0001).

### Effects of Extra Motivation on Brain Activity

Compared with standard trials, extra motivation trials had higher ΔHbO_2_ in all brain regions tested (Fig 2, Table 3). The magnitude of this extra motivation effect was greater for SFG9m than every other brain region (p≤0.0386).

**Figure 2.**
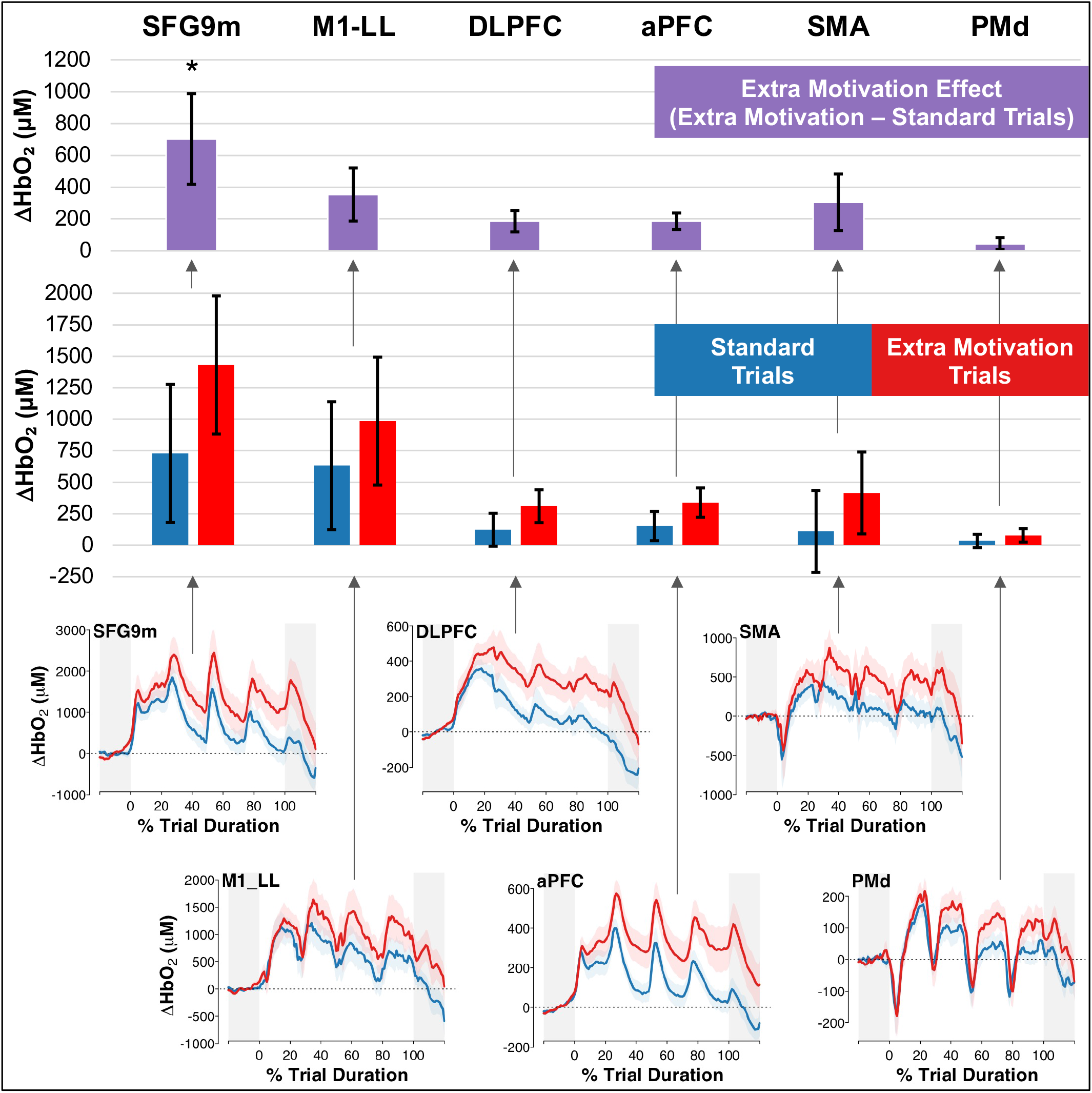
Oxygenated hemoglobin responses (ΔHbO_2_) by task condition and brain region. Bars show mean estimates and error bars show 95% CI for ΔHbO_2_ from baseline, controlled for trial heart rate. Bottom panels show the ΔHbO_2_ time series averaged across trials and participants for each task condition, with a shaded region to depict the standard error across participants. *The extra motivation effect was significantly greater for SFG9m than every other region of interest. SFG9m, superior frontal gyrus medial area 9; M1-LL, primary motor cortex lower limb area; DLPFC, dorsolateral prefrontal cortex; aPFC, anterior prefrontal cortex; SMA, supplementary motor area; PMd, dorsal premotor cortex

**Table 3.** Oxygenated hemoglobin responses (ΔHbO_2_) by task condition and brain region1.

| **Brain Region** | **Standard Trials** | **Extra Motivation Trials** | **Extra Motivation Effect**<br>**(Extra – Standard)** |
| --- | --- | --- | --- |
| **SFG9m** | 728 [180, 1277] | 1431 [883, 1979] | 703 [418, 988]* |
| **M1-LL** | 631 [124, 1139] | 985 [478, 1493] | 354 [187, 521] |
| **DLPFC** | 123 [-7, 254] | 309 [179, 440] | 186 [119, 253] |
| **aPFC** | 152 [36, 269] | 338 [222, 455] | 186 [134, 238] |
| **SMA** | 110 [-215, 435] | 415 [90, 740] | 305 [127, 483] |
| **PMd** | 33 [-20, 87] | 78 [25, 132] | 45 [7, 83] |
| Values are mean estimates [95% CI] for $\Delta\text{HbO}_2$ from baseline in $\mu\text{M}$, controlled for trial heart rate. *The extra motivation effect was significantly greater for SFG9m than every other region of interest. SFG9m, superior frontal gyrus medial area 9; M1-LL, primary motor cortex lower limb area; DLPFC, dorsolateral prefrontal cortex; aPFC, anterior prefrontal cortex; SMA, supplementary motor area; PMd, dorsal premotor cortex |  |  |  |

### Brain Activity Correlations with Self-Reported Exercise Motivation Across Participants

Across participants, more self-determined motivation for exercise was correlated with greater mean brain ΔHbO_2_ responses to extra motivation (relative to standard trials) in SFG9m and aPFC (Table 4). None of the individual BREQ motivation domains contributing to the overall self-determination score showed significant correlations with the brain ΔHbO_2_ responses, but the estimated associations for SFG9m and aPFC were all positive for autonomous forms of motivation and negative for controlled forms.

**Table 4.** Behavioral associations with brain oxygenated hemoglobin responses (ΔHbO_2_)

|  | Brain Region |  |  |  |  |  |
| --- | --- | --- | --- | --- | --- | --- |
|  | SFG9m | M1-LL | DLPFC | aPFC | SMA | PMd |
| **Self-reported motivation for exercise (BREQ score) associations with mean ΔHbO<sub>2</sub> response to extra motivation** (relative to standard trials). Partial correlations, controlled for trial heart rate. |  |  |  |  |  |  |
| **Overall self-determined motivation**<br>(Autonomous – Controlled) | 0.51* | 0.25 | 0.37 | 0.47* | 0.00 | 0.14 |
| **Autonomous motivation**<br>(Intrinsic + Identified) | 0.32 | 0.26 | 0.34 | 0.32 | 0.02 | 0.28 |
| **Intrinsic motivation** | 0.28 | 0.23 | 0.36 | 0.23 | 0.08 | 0.19 |
| **Identified motivation** | 0.31 | 0.26 | 0.29 | 0.35 | -0.03 | 0.32 |
| **Controlled motivation**<br>(Introjected + External) | -0.23 | 0.01 | -0.03 | -0.19 | 0.02 | 0.17 |
| **Introjected motivation** | -0.26 | -0.06 | -0.02 | -0.24 | 0.00 | 0.02 |
| **External motivation** | -0.14 | 0.08 | -0.04 | -0.09 | 0.04 | 0.25 |
| **Trial-by-trial ΔHbO<sub>2</sub> associations with walking speed.** Partial repeated-measures correlations (within-participants), controlled for trial type (extra motivation vs. standard). |  |  |  |  |  |  |
| **Walking speed** | 0.19* | 0.09 | 0.07 | 0.06 | 0.16* | 0.22* |
| *p<0.05. SFG9m, superior frontal gyrus medial area 9; M1-LL, primary motor cortex lower limb area; DLPFC, dorsolateral prefrontal cortex; aPFC, anterior prefrontal cortex; SMA, supplementary motor area; PMd, dorsal premotor cortex; BREQ, Behavioral Regulation for Exercise Questionnaire |  |  |  |  |  |  |

### Brain Activity Correlations with Walking Speed Across Trials

Within participants and conditions (extra motivation vs. standard), greater brain ΔHbO_2_ increases in SFG9m, SMA and PMd were correlated with faster walking speed across trials (Table 4).

## Discussion

This walking exercise study among non-disabled adults experimentally assessed brain activity changes during periods of additional motivation, and examined how this brain activity relates to self-reported motivation and walking speed. The exercise protocol appeared to provide sufficient motivational challenge since it required vigorous aerobic exertion. Faster walking during trials randomized to include ‘extra motivation’ also showed that the experimental manipulation successfully altered participant behavior. As hypothesized, SFG9m was more active during ‘extra motivation’ trials and more responsive to the ‘extra motivation’ condition than all other brain regions tested. Further, greater SFG9m activity was associated with more self-determined motivation for exercise and faster walking speeds. These results substantially strengthen prior findings from correlational MRI studies,^28, 29^ indicating that SFG9m plays an important role in motivation for walking activity. This brain region has received little previous attention, but now appears to be a promising target for future neuromodulation studies and prognostic research aiming to augment physical activity motivation or predict walking-related outcomes (e.g. for gait rehabilitation).

Across individuals, greater SFG9m (and aPFC) response to the ‘extra motivation’ condition was specifically associated with more *self-determined* motivation for exercise, suggesting that these regions may specifically regulate more autonomous and less controlled behavior. More autonomous or self-determined exercise motivation is generally considered to have more positive effects on exercise behavior and experiences,^12^ which implies beneficial effects of higher SFG9m activity. Brain activity correlations with the different domains of self-reported motivation may also reflect the extent to which the experimental manipulation targeted each motivation domain. From this perspective, it may initially seem counterintuitive that the additional *external inputs* (e.g. lap timer, verbal encouragement) in the ‘extra motivation’ trials appeared to be more effective at upregulating brain activity for individuals who reported more *intrinsic* (and less *extrinsic*) forms of *motivation* for exercise.

In the broader psychology literature, it was originally proposed that external rewards could only generate extrinsic motivation,^51^ and would diminish intrinsic motivation by reducing perceived autonomy.^52^ However, general interest theory^53^ subsequently departed from this former view (cf.^51^) based on evidence that external rewards can actually enhance intrinsic motivation under more natural experimental conditions.^54, 55^ For example, external rewards that are contingent on meeting a specific, challenging and attainable performance criterion appear to increase perceived autonomy and competence, which can generate intrinsic motivation.^53, 56^ The gain in perceived autonomy with this type of reward contingency is thought to occur because knowing what level of performance is needed to receive a reward gives the individual more control over their own reinforcement and environment.^53, 56^ Thus, our display of the participant’s best lap time and positive feedback for each new record may have been important stimuli for generating more autonomous forms of motivation.Future studies could benefit gait rehabilitation and exercise interventions by more specifically identifying inputs and contextual conditions that optimize self-determined engagement in walking exercise.

The positive association we observed between brain responses and self-determined motivation was based on differences between individuals. Therefore, it is also possible that our external motivational inputs may have had more intrinsic salience to some participants. Individual differences in interests and experiences can influence the extent to which a particular activity is intrinsically motivating.^57^ Additional factors (e.g. physical and social environment) may also contribute to an individual’s motivation for physical activity.^58^ These potential individual differences should be considered when aiming to augment motivation in future studies, exercise programs and clinical rehabilitation.

In addition to these between-participant differences, we also analyzed within-participant differences across trials and found that greater SFG9m, SMA and PMd activity was associated with faster walking speed, while controlling for the effects of trial condition (extra motivation vs. standard). Unlike SMA and PMd, SFG9m is not typically considered a motor region, but this finding adds to a growing body of evidence that SFG9m may indeed be a novel motor area, or at least highly relevant to walking function.^28, 29, 59, 60^ For example, the medial prefrontal cortex (including SFG9m) was recently found to be a major origin of a key descending projection for gross motor functions like walking (the cortico-reticulo-spinal pathway).^59, 60^

The current study cannot determine whether SFG9m exerted any direct effects on walking through the cortico-reticulo-spinal pathway and/or indirect effects via connections with established motor regions of the cortex (M1-LL, SMA, PMd). Given the correlational nature of this result, it is also possible that SFG9m did not exert any effects on walking (non-causal/confounded association) or that faster walking somehow led to greater SFG9m activation (reverse causality). However, it is interesting that SFG9m was the only tested region that was associated with both self-reported motivation and walking speed, suggesting that it could be a key region linking affective signaling to motor output.

It is also noteworthy that all brain regions tested were more active during the ‘extra motivation’ condition than the standard trials, including SFG9m, M1-LL, SMA, PMd, aPFC and DLPFC. These findings indicate that simple methods to augment motivation during walking exercise (e.g. timing trials, providing feedback about speed and encouragement to beat previous records) can upregulate brain activity across the motor cortex and beyond. Similar motivational techniques have demonstrated feasibility among clinical populations with gait impairment,^20, 61^ and have shown efficacy for improving gait outcomes.^20^ The current findings provide a possible neurobiological explanation for *how* this motivational input during rehabilitation might improve outcomes (via widespread upregulation of brain activity).

### Limitations

This study attempted the challenging task of elucidating the neural underpinnings of a complex psychological construct (motivation), for which there is no flawless approach.^62-64^ While our randomized task comparison design is widely used (e.g. in the fMRI literature) and has important strengths over other methods, it still does not guarantee valid causal inference.^63^ This design evaluates the effects of brain activity on behavior by manipulating the dependent variable (behavior -> brain activity), which assumes a 1:1 relationship between the two.^63^ When evaluating the brain activity underlying motivation, this strong assumption may be particularly tenuous because it is difficult to design experimental conditions that specifically alter motivation without also affecting other neural processes. For example, auditory stimulation was also likely higher in our ‘extra motivation’ condition (versus standard trials) and cognitive processing may also have been more demanding because of the additional stimuli. Addressing the uncertain causal inference from our current results will likely require additional studies targeting the same question with complementary designs.^62, 63^

As referenced above, the analyses correlating brain activity with self-reported motivation and walking speed may have also been confounded by other brain or behavioral variables. We attempted to mitigate this issue by statistically controlling for the most plausible confounding variable in each analysis, but false positive or negative findings are still possible. With our moderate sample size (N=26), the large number of repeated trials provided sufficient power to compare brain activity between conditions and to detect even relatively small brain activity associations with walking speed between trials. However, we may have been underpowered to detect smaller brain activity associations with self-reported motivation between participants. Mobile brain imaging technology like fNIRS is also limited to measuring activity from more superficial brain structures and we did not obtain MRI data in this study to confirm that fNIRS recordings were made from the same brain locations across participants.

## Conclusions

For walking exercise among non-disabled adults, clinically-relevant inputs designed to increase motivation (e.g. timing trials, providing feedback about speed and encouragement to beat previous records) elicit widespread increases in cerebral cortex activity. These increases were the greatest in SFG9m, a region of the medial prefrontal cortex with recently-discovered relevance to motor function. Greater activity increases in SFG9m were also related to more self-determined motivation for exercise and faster walking speed, suggesting that it may play a key role in motivation for physical activity.**Trials**

## Notes

**Funding:** This study was supported by the National Institutes of Health (R01HD093694 and UL1TR001425) and the Oliver Family Foundation.

**Conflicts of Interests:** The authors declare no conflicts of interest.

### Competing Interest Statement

The authors have declared no competing interest.

